# RhoGEF9 splice isoforms influence neuronal maturation and synapse formation downstream of *α*2 GABA_A_ receptors

**DOI:** 10.1101/197905

**Authors:** Claire deGroot, Amalia Floriou-Servou, Yuan-Chen Tsai, Simon Früh, Manuela Kohler, Georgia Parkin, Cornelia Schwerdel, Giovanna Bosshard, Kai Kaila, Jean-Marc Fritschy, Shiva K. Tyagarajan

## Abstract

In developing brain neuronal migration, dendrite outgrowth and dendritic spine outgrowth are controlled by Cdc42, a small GTPase of the Rho family, and its activators. Cdc42 function in promoting actin polymerization is crucial for glutamatergic synapse regulation. Here, we focus on GABAergic synapse-specific activator of Cdc42, collybistin (CB) and examine functional differences between its splice isoforms CB1 and CB2. We report that CB1 and CB2 differentially regulate GABAergic synapse formation *in vitro* along proximal-distal axis and adult-born neuron maturation *in vivo*. The functional specialization between CB1 and CB2 isoforms arises from their differential protein half-life, in turn regulated by ubiquitin conjugation of the unique CB1 C-terminus. We report that CB1 and CB2 negatively regulate Cdc42; however, Cdc42 activation is dependent on CB interaction with gephyrin. During hippocampal adult neurogenesis CB1 regulates neuronal migration, while CB2 is essential for dendrite outgrowth. Finally, using mice lacking *Gabra2* subunit, we show that CB1 function is downstream of GABA_A_Rs, and we can rescue adult neurogenesis deficit observed in *Gabra2* KO. Overall, our results uncover previously unexpected role for CB isoforms downstream of α2-containing GABA_A_Rs during neuron maturation in a Cdc42 dependent mechanism.

**Author Summary:** GABAergic inhibition regulates distinct stages of brain development; however, cellular mechanisms downstream of GABA_A_ receptors (GABA_A_Rs) that influence neuronal migration, maturation and synaptogenesis are less clear. *ArfGEF9* encodes for RhoGEF with Cdc42 and TC10 GTPase as its substrates. Interestingly, ArhGEF9 is the only known RhoGEF essential for GABAergic synapse formation and maintenance. We report that during brain development ArfGEF9 mRNA splicing regulation generates different ratios of CB1 and CB2 splice isoforms. CB1 mRNA splicing is enhanced during early brain developmental, while CB2 levels remains constant throughout brain development. We also show that CB1 protein has shorter half-life and ubiquitin proteasome system restricts CB1 abundance within developing neuron to modulate neuron migration and distributing GABAergic synapse along the proximal-distal axis. On the other hand, CB2 isoform although expressed abundantly throughout brain development is essential for neuron dendrite maturation. Together, our data identifies specific post-transcriptional and post-translational mechanisms downstream of GABA_A_Rs influencing ArhGEF9 function to regulate distinct aspects of neuronal maturation process.

## Introduction

During CNS development, GABAergic transmission regulates key steps of neurogenesis and neuronal circuit formation [1,2]. A similar role has been observed also for the regulation of adult neurogenesis [3-5]. The diverse effects of GABA can be observed best upon inactivation of specific GABA_A_R subtypes by targeted deletion of the a subunit variants, impairing cell migration, dendrite formation and synaptic integration [6,7]. Furthermore, functional inactivation of the scaffolding protein gephyrin, present at GABAergic synapses, strongly impairs formation and growth of dendrites, presumably by reducing GABAergic transmission [8]. Hence, gephyrin and other postsynaptic proteins represent essential components of downstream signaling and cytoskeletal function associated with GABAergic synapses [7].

In addition to GABA_A_R and gephyrin, several proteins have been found to be essential for GABAergic synapse structure and function [9]. Among these, collybistin (CB) is of special interest, as it has emerged as a key organizer of GABAergic synapses. CB is a member of the Dbl family of guanine nucleotide exchange factors (GEF) that, through interaction with gephyrin and the postsynaptic cell adhesion protein neuroligin 2, contributes to the formation and stabilization of GABAergic synapses [10-12]. CB selectively activates Cdc42, albeit weakly [13] and TC10, but their roles down-stream of CB remains enigmatic. Targeted deletion of Cdc42 has been reported not to affect gephyrin clustering [14], whereas, *in vitro*, Cdc42 overexpression can rescue gephyrin clustering impaired by the presence of a CB mutant lacking the pleckstrin homology (PH) domain [15]. In addition, Cdc42, either alone or in combination with CB, modulates the size and shape of postsynaptic gephyrin clusters [15]. A major unresolved issue about CB is the functional role of alternative splicing of its mRNA, giving rise to three main isoforms differing exclusively in the C-terminal domain (CB1-3) and having or lacking an SH3 domain close to the N-terminus. While the latter is thought to regulate the catalytic activity of CB, the specific functions of the CB1-3 isoforms are currently not understood [16]. CB3 is the ortholog of human hPEM2, mutations of which are associated to two cases of X-linked mental retardation without hyperekplexia, associated with epilepsy, anxiety, and sensory hyperarousal [17,18]. CB1, CB2 and CB3 have previously been shown to exert comparable effects on postsynaptic gephyrin clustering when overexpressed in primary neurons [19].

In this study, we investigated possible functional differences between CB isoforms on neuronal development and GABAergic synapse formation in vitro and in vivo, using transfection of primary neurons as well as neuronal precursor cells of the dentate gyrus; and we characterized possible isoform-specific properties of CB1 and CB2 and their interactions with Cdc42 using biochemical assays. Based on previous observations that, upon overexpression, CB isoforms are not restricted to GABAergic postsynaptic sites but distributed in the soma and entire dendritic tree [15], we reasoned that CB might have distinct functions in synapses and on the cytoskeleton, which might depend also on interaction with gephyrin and/or GABA_A_R. Our results demonstrate that CB1 and CB2 differ in their regulation of GABAergic synapse formation and dendritic growth in developing neurons. Unique lysine residues in the C-terminus of CB1 determine its short half-life compared to CB2 and explain major functional differences between the two isoforms. Further, we show that CB interaction with Cdc42 regulates its action on the cytoskeleton and is modulated by gephyrin. Paradoxically, silencing CB expression exacerbates Cdc42 function, providing a basis for functional differences between CB1 and CB2 that depend on their differential half-life. Finally, by using targeted deletion of the GABA_A_R α2 subunit, we demonstrate that CB isoforms act downstream of α2 GABA_A_R in adult-born dentate gyrus granule cells.

## Results

### CB1 and CB2 isoforms exhibit differential effects on gephyrin clustering *in vitro*

We aim here to understand the functional significance of CB1 and CB2 isoforms for GABAergic synapse formation. CB1 and CB2 isoforms differ primarily in their C-terminus (93% overall sequence identity; unique 30 amino acid and 8 amino acid C-terminus, respectively), which suggests a specific role for this region of the protein in setting isoform-specific properties. Given the complexity in specifically targeting each of the CB isoforms (CB1_SH3+_, CB1_SH3-_, CB2_SH3+_ and CB2_SH3-_) individually *in vivo*, we resorted to an overexpression system to individually express these four cDNAs for dissecting out their functional differences. We cloned CB1 and CB2 isoforms from a rat cDNA library and added either eGFP, mCherry or V5 N-terminal tags (Fig. 1A). eGFP-gephyrin was co-transfected with either of the mCherry-CB isoforms into primary hippocampal neurons at 8 days in vitro (DIV) to facilitate visualization of GABAergic synapses, and analyzed for changes in density of postsynaptic gephyrin clusters 7 days post-transfection (DIV 8 + 7). Representative images of neurons expressing either eGFP-gephyrin alone, or together with mCherry-CB (blue) isoforms opposed to vesicular GABA transporter (vGAT)-positive terminals (red) are shown (Fig. 1B-F). Independent studies have demonstrated that alterations in postsynaptic gephyrin scaffolds are reflected by corresponding changes in GABAergic transmission [20-22]. Here, we confirmed that eGFP-gephyrin clusters along the proximal-distal axis of dendrites are associated with GABA_A_Rs by co-labeling for the γ2 GABA_A_R subunit (Fig. 1B-F, lower panel).

**Figure 1:**
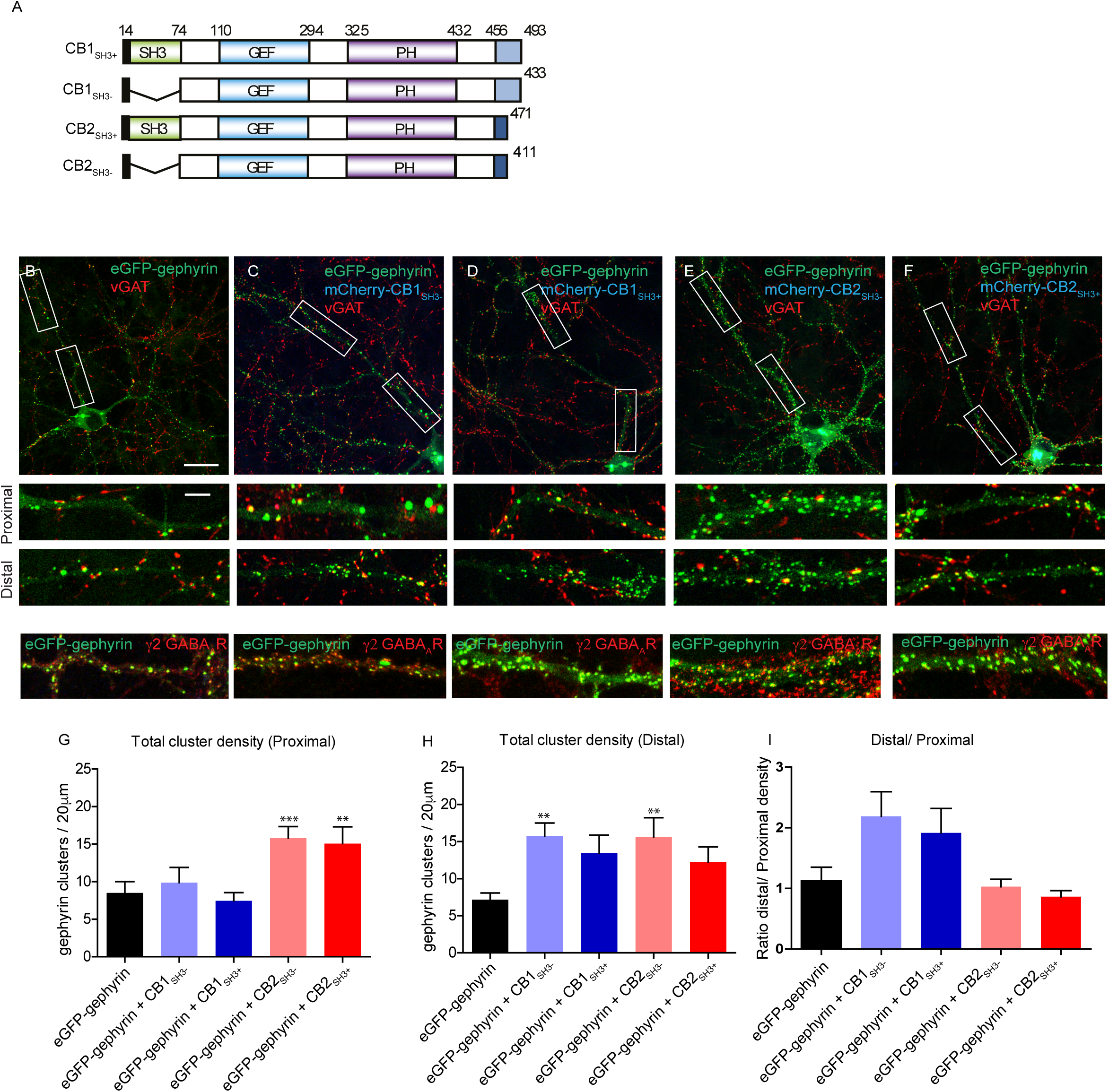
V5-CB1 splice isoforms enhance eGFP-gephyrin clustering along proximal-distal axis. (A) Cartoon of various CB isoforms and deletion mutations used in the current study. **(B-F)** 8 + 7 DIV neurons co-transfected with either V5-CB1_SH3+_, V5-CB1_SH3-_, V5-CB2_SH3+_ or V5-CB2_SH3-_ isoform (blue) and eGFP-gephyrin (green) and stained for presynaptic marker VGAT (red). Alterations in the morphology of eGFP-gephyrin synaptic clusters was observed. **(B-F, lower panels)** zoom of dendritic proximal or distal dendritic segment showing eGFP-gephyrin clustering and vGAT co-localization. The localization of eGFP-gephyrin clusters with γ 2 GABA_A_R subunit was also observed. **(G-I)** Quantification of eGFP-gephyrin synaptic cluster density per 20 μm dendrite (DIV 8 + 7) at proximal and distal dendritic segments of neurons co-expressing eGFP-gephyrin and mCherry-CB1/2 isoforms. Scale bar 10μm and 5 μm. Statistical analysis for cluster density; One-way ANOVA, Bonferonni post-hoc test, p = 0.85; size analysis: Kruskal-Wallis non parametric test, Dunn’s multiple comparison test p<0.0001.

Examination of dendritic proximal segments (up to 40 μm from soma; upper panels) and distal segments (80-120 μm from soma; lower panels) revealed a differential distribution of eGFP-gephyrin clusters selectively in the presence of CB1_SH3+_ or CB1_SH3-_ (Fig. 1G-I). Quantification showed that CB1_SH3-_ and CB1_SH3+_ specifically enhance eGFP-gephyrin cluster density in distal dendritic segments, while CB2_SH3-_ and CB2_SH3+_ enhance eGFP-gephyrin clustering uniformly throughout the entire dendrites (Fig. 1G-I; Table 1). These results demonstrate that CB splice isoforms differentially influence gephyrin clustering (i.e., GABAergic synapse formation) in primary neurons in a compartment-specific manner, pointing towards a functional segregation of CB isoforms. *In vivo*, proximal and distal dendrites of pyramidal cells are targeted by different interneurons; as this compartment-specific organization is not preserved *in vitro*, we presume that CB1 mainly facilitates synapse formation in immature segments of growing dendrites.

**Table 1.** Quantitative analysis of eGFP-gephyrin cluster density in primary neurons. The density is represented per 20 μm dendrite length. Proximal dendrite; One-way ANOVA (F 4.38, 4) = 26; Distal dendrite; One-way ANOVA (F 3.4, 4) = 26. Bonferroni post-hoc test for pair wise comparison.

| Condition | Cluster density (proximal) | 1 way ANOVA | Cluster density (distal) | 1 way ANOVA |
| --- | --- | --- | --- | --- |
| eGFP-gephyrin | 6.9 ± 0.9 |  | 6.7 ± 0.6 |  |
| eGFP-gephyrin + mCherryCB1<sub>SH3-</sub> | 8.8 ± 1.3 | P<0.0001 | 15.2 ± 1.5 | P=0.0012 |
| eGFP-gephyrin+mCherry-CB1<sub>SH3+</sub> | 7.6 ± 0.7 | P<0.0001 | 12.7 ± 1.5 | P=0.0012 |
| eGFP-gephyrin + mCherryCB2<sub>SH3-</sub> | 14.3 ± 1.3 | P<0.0001 | 15.1 ± 1.7 | P=0.0012 |
| eGFP-gephyrin+mCherry-CB2<sub>SH3+</sub> | 15.1 ± 1.4 | P<0.0001 | 12.1 ± 1.8 | P=0.0012 |
| **Ub mutants** |  |  |  |  |
| eGFP-gephyrin | 6.9 ± 1 |  | 6.5 ± 0.7 |  |
| eGFP-gephyrin + V5CB1<sub>SH3-</sub> | 7.9 ± 0.7 | P<0.0001 | 13.6 ± 0.9 | P<0.0001 |
| eGFP-gephyrin+V5-CB1<sub>SH3+</sub> | 6.8 ± 1 | P<0.0001 | 11.4 ± 1.2 | P<0.05 |
| eGFP-gephyrin + V5CB1<sub>SH3-</sub> 2R | 18.9 ± 1.1 | P<0.0001 | 13.8 ± 0.8 | P<0.0001 |
| eGFP-gephyrin+mCherry-CB1<sub>SH3+</sub> 2R | 20.6 ± 1.6 | P<0.0001 | 16.1 ± 1.7 | P<0.0001 |

### CB1 isoform is spliced differentially during brain development

As a corollary, we reasoned that for CB1 to act in immature neurons, it should be expressed early during brain development. To test this possibility, we collected total mRNA from rat neocortex, hippocampus and cerebellum at different time-points after birth (P5, P9, P15, P30) and performed quantitative real-time PCR (qRT-PCR) analysis of CB1, CB2 and PanCB transcript levels. Normalizing the samples to PanCB levels allowed us to evaluate the relative ratios of CB1 and CB2 mRNA (Fig. 2A-C). In all samples, CB2 levels were similar to panCB, confirming that it represents the main splice isoform in the CNS across development (One-way ANOVA F (3, 16) = 3.9; P = 0.02). Nevertheless, the level of CB1 mRNA was consistently higher at the early time-points in the three brain regions tested with a gradual reduction over time. Interestingly, CB1 mRNA in the neocortex even represented almost 60% of the total CB at P5, and decreased to 18% at P30 (One-way ANOVA F (3, 16) = 299; P<0.0001). In contrast, hippocampus and cerebellum samples showed 20% total CB1 mRNA at P5, reducing to below 10% of PanCB by P30 (One-way ANOVA F (3,16) = 4.2; P = 0.0219; One-way ANOVA F (3,16) = 10.5; P = 0.0005). Thus, CB1 might play a predominant role during the early phase of postnatal neuronal maturation, when rapid changes in dendritic growth and synapse formation are taking place.

**Figure 2:**
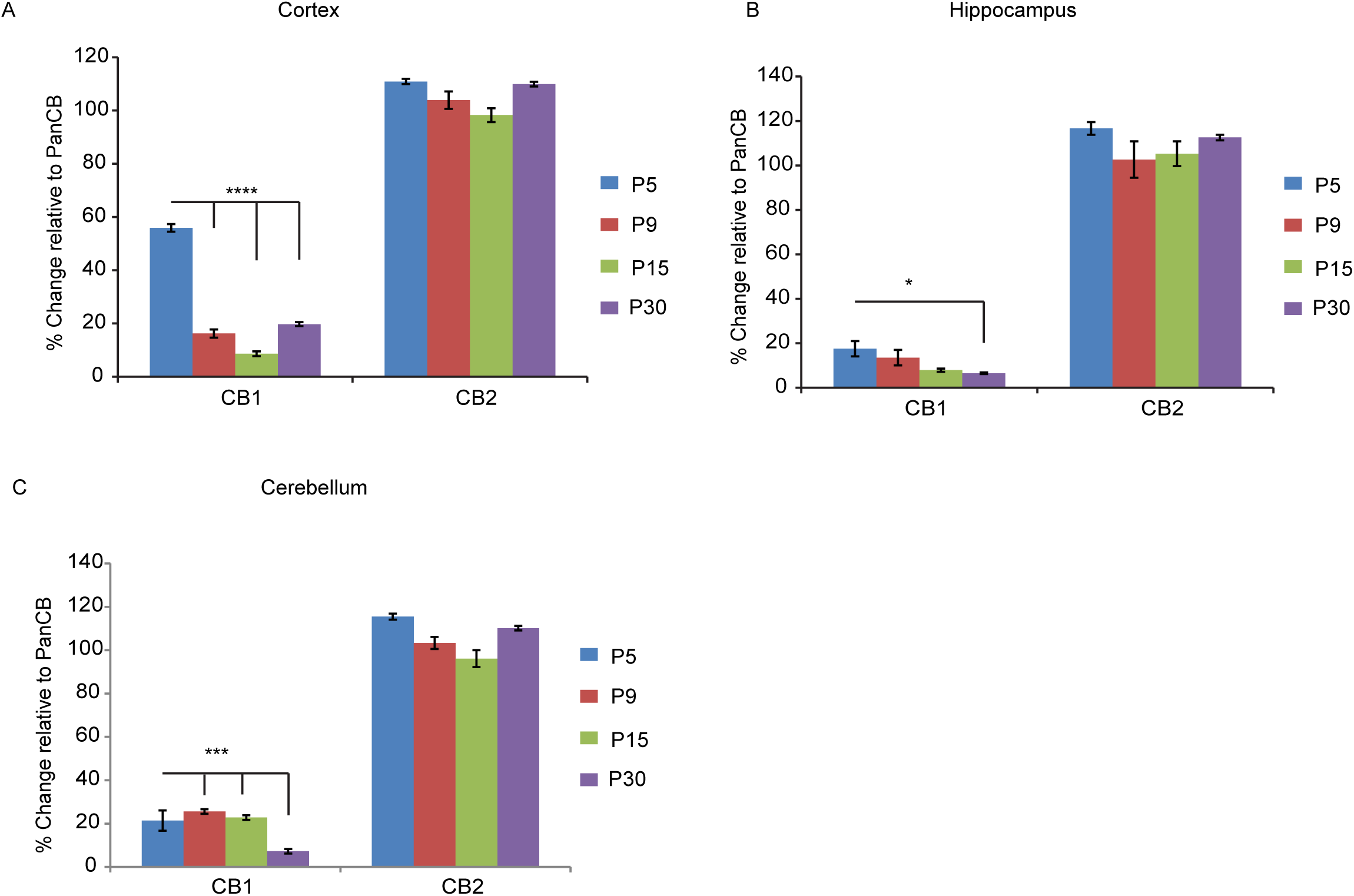
CB1 mRNA alternative splicing is altered during brain development. (A) CB1 and CB2 mRNA expression levels in the cortex at P5, P9, P15 and P30 developmental stages normalized to PanCB mRNA levels. **(B)** CB1 and CB2 mRNA expression levels in the hippocampal formation at P5, P9, P15 and P30 developmental stages normalized to PanCB mRNA levels. **(C)** CB1 and CB2 mRNA expression levels in the cerebellum at P5, P9, P15 and P30 developmental stages normalized to PanCB mRNA levels.

### CB isoforms differentially influence specific stages of adult neurogenesis

To explore the functional specificity of CB isoforms *in vivo*, we moved to adult neurogenesis in the subgranular zone (SGZ), which gives rise to new dentate gyrus granule cells (GCs) [23]. This model provides an amenable *in vivo* system to test in adult mice the influence of signaling downstream of GABA_A_R on neuronal maturation in a cell-autonomous manner.

We used retroviruses encoding eGFP and eGFP-tagged CB1 and CB2 isoforms (see materials and methods) to infect dividing neural progenitor cells in 8-10 week-old mice. Successful labeling of adult-born neurons allowed us to follow their position in the granule cell layer and morphology over a long time span. We first compared the effects of eGFP-CB1_SH3-_ or eGFP expression in adult-born neurons, focusing on neuronal migration and dendrite maturation at 14, 28 or 42 days post-virus injection (dpi). These time-points represent three distinct phases of maturation of adult-born GCs and allow direct comparison with earlier studies [6].

GCs overexpressing CB1_SH3-_ penetrated less deeply into the GCL, with 80-90% of cells remaining within <20 μm from the SGZ border at each of the three time-points examined, whereas in the control group, >25% of GCs migrated more than 20 μm. The difference was significant at 28 and 42 dpi (Kolmogorov-Smirnov test; P <0.05; Fig. 3A-A’’), suggesting that CB1_SH3-_ negatively regulates cytoskeletal reorganization required for cell motility. However, all transduced GCs had moved away from the SGZ, indicating that migration *per se* was not completely impaired. We then tested the influence of eGFP-CB2_SH3-_ overexpression during neurogenesis in an independent batch of mice. Analysis of eGFP-CB2_SH3-_ infected neurons at 14, 28 and 42 dpi showed no significant migration differences compared to control (Kolmogorov-Smirnov test; P = 0.438, 0.566 and 0.89 respectively; Fig. 3B-B’’).

**Figure 3:**
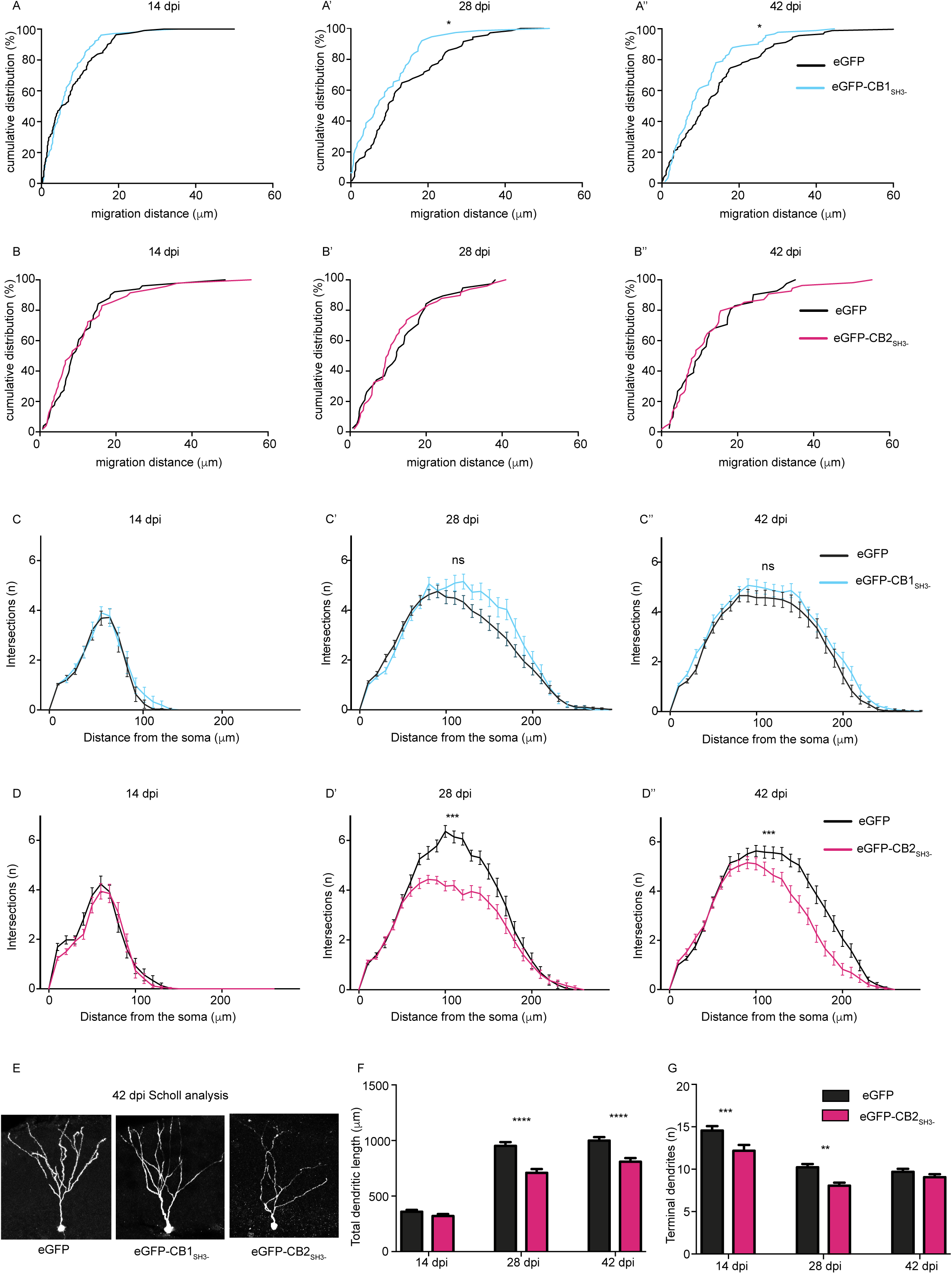
CB1 and CB2 splice isoforms influence adult neurogenesis. (A-A”) WT mice infected with eGFP or eGFP-CB1_SH3-_ retrovirus; **(B-B”)** WT mice infected with eGFP or eGFP-CB2_SH3-_ retrovirus and analyzed for cell migration at 14 dpi, 28dpi and 42 dpi. **(C-C”)** WT mice infected with eGFP or eGFP-CB1_SH3-_ retrovirus; **(D-D”)** WT mice infected with eGFP or eGFP-CB2_SH3-_ retrovirus and analyzed for dendrite maturation using Sholl analysis at 14 dpi, 28dpi and 42 dpi. **(E)** Morphology of adult newborn neurons infected with retrovirus expressing eGFP, eGFP-CB1_SH3-_ or eGFPCB2_SH3-_. **(F)** Total dendritic length of newborn neurons infected with eGFP, eGFPCB1_SH3-_ or eGFP-CB2_SH3-_. **(G)** Total number of terminal dendrites expressing eGFP or eGFP-CB2_SH3-_. Kolmogorov-Smirnov (A-A”, B-B”), N = 38 to 54 cells/group. Mann-Whitney test for AUC (C-C”, D-D”), N = 31-65 cells/group. Two-way ANOVA, Bonferroni post-hoc test (F,G), N = 34-45 cells/group.

Next, we quantified dendritic complexity by Sholl analysis, which likely reflects on neuronal maturation and cytoskeleton regulation. Overexpression of eGFP-CB1_SH3-_ did not significantly influence dendritic arborization compared to eGFP-control neurons (Mann Whitney test for AUC; P = 0.7712, 0.0979, 0.0852 respectively; Fig. 3C-C’’). However, eGFP-CB2_SH3-_ overexpression impaired arborization, as seen by the significant reduction in complexity of dendritic tree compared to eGFP-cells (Mann Whitney test for AUC; P = 0.1714 for D; P<0.0001 for D’, P<0.0001 for D’’; Fig. 3D-D”).

To confirm this observation, we also determined the total dendritic length of eGFP, eGFP-CB1_SH3-_ and eGFP-CB2_SH3-_ infected adult newborn neurons (Fig. 3E). eGFP-CB2_SH3-_ overexpression reduced total dendritic length and terminal dendrite length (Fig. 3F-G). Furthermore, eGFP-CB2_SH3-_ overexpression caused a significant effect of time (F (2, 66) = 52.70, P = 0.0001) compared to eGFP control (F (1, 66) = 10.24, P = 0.021). These observations indicate that CB1_SH3-_ and CB2_SH3-_ isoforms regulate distinct processes during adult newborn neuron maturation, despite overexpression and the presence of endogenous CB isoforms.

### CB1 has shorter protein half-life than CB2.

In order to elucidate the mechanistic basis of these functional differences, we turned to protein biochemistry. We analyzed the protein half-life of CB1 and CB2 isoforms in HEK-293T cells, based on the fact that the C-terminus of CB1 and CB2 contain different lysine residues, which might be targeted by ubiquitination [24]. To achieve this goal, we blocked protein synthesis using cyclohexamide (100 μM) 12 h post-V5-CB isoform transfection and performed Western blotting (WB) for V5 to measure the relative abundance of CB isoforms at different time points (Fig. 4A-D). This analysis showed that both CB1_SH3+_ and CB1_SH3-_ have significantly shorter half-life than CB2_SH3+_ or CB2_SH3-_. Interestingly, CB1_SH3-_ exhibited the shortest half-life (1.8 hr) and CB2_SH3-_ was the most stable of the four isoforms tested (7.3 hr). We further determined the CB1_SH3-_ protein half-life in primary hippocampal neurons infected with lentiviruses encoding mCherry-CB1_SH3-_ and we obtained a very similar result (Fig. 4E), suggesting that CB protein regulation might be conserved between cell types.

**Figure 4:**
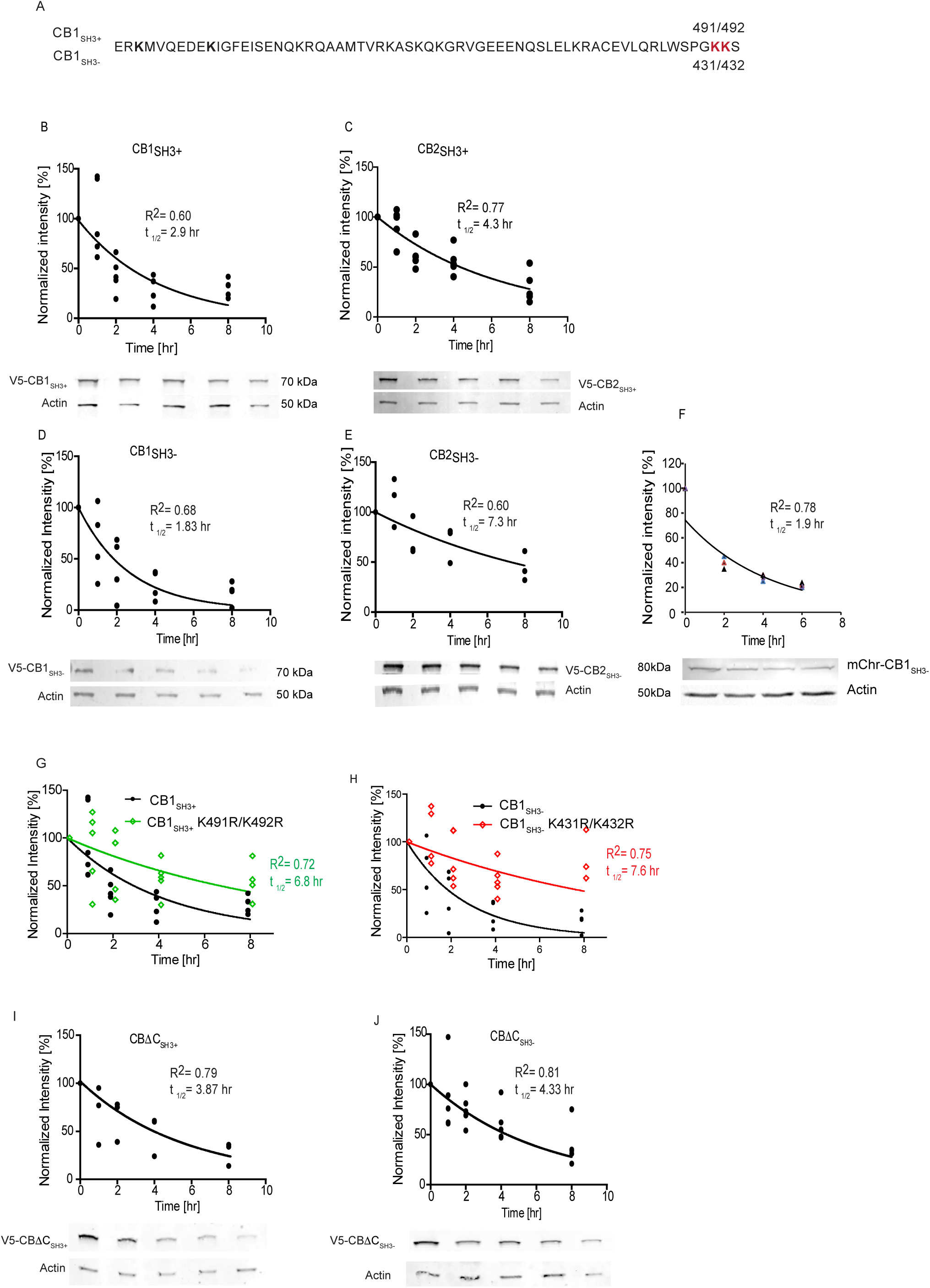
Protein half-life of V5-CB1 and V5-CB2 splice isoforms differ. (A) Protein stability of V5-CB1_SH3+_ was determined by collecting the protein samples at 0, 1, 2, 4 and 8 hours in the presence of cyclohexamide (CHX) (0.1mM) and half-life relative to actin was plotted. The t_1/2_ and R^2^ values were determined (inset). **(B)** Protein half-life of CB2_SH3+_ isoform. **(C)** Protein half-life of CB1_SH3-_ in HEK-293T cells. **(D)** Protein half-life of CB2_SH3-_ isoform in HEK-293T cells. CB1_SH3-_ shows the shortest half-life (1.8 hr), while CB2_SH3-_ has the longest half-life (7.294 hr). **(E)** Protein half-life of mCherry-CB1_SH3-_ in primary hippocampal neuron is similar to HEK-293T cells. **(F)** CB1 isoform C-terminal amino acid sequence with specific Ub lysine residues marked in red. **(G-J)** The predicted K491R/K492R ubiquitin residues in CB1 were mutated Lys/Arg and the protein stability of the mutants was determined. The ubiquitin site mutants have a longer half-life compared to the WT counterparts. **(I-J)** CBΔC_SH3+_ and CBΔC_SH3-_ C-terminus deletion mutations show similar protein half-life.

We then checked whether CB1_SH3-_ and CB2_SH3-_ are substrates for ubiquitination. We treated one set of HEK-293T cells with the proteasomal inhibitor MG132 or vehicle (DMSO) and examined for HA-Ubiquitin (Ub) conjugation of V5-CB isoforms (Suppl. Fig. 1A). Immunoprecipitation for HA, followed by WB for V5 showed higher Ub conjugation in MG132-treated samples in comparison to control. We confirmed this observation by performing the experiment in reverse. We immunoprecipitated V5-CB and probed for HA-Ub conjugation by performing WB against HA. We could see a distinct increase in HA-Ub conjugation of all V5-CB isoforms in the presence of MG132 (Suppl. Fig. 1B).

CB1 harbors two lysine residues in the penultimate position, suggesting that they might regulate its protein stability. We mutated these two lysine residues to arginine in both CB1_SH3+_ (491/492) and CB1_SH3-_ (431/432) isoforms (Fig. 4F) and determined the half-life of these CB1 lysine mutants in HEK-293T cells as before. We found a significantly increased protein stability for both isoforms (Fig. 4G-H); for instance, mutation of lysine (431/432) in CB1_SH3-_ increased the protein half-life to similar levels as seen for CB2_SH3-_ (7.6 hr), suggesting that these lysine residues on CB1 largely determine protein turnover. The global contribution of C-terminus sequence in regulating protein stability of CB isoforms is still not fully elucidated. Hence, we deleted the unique C-terminus of CB1 to remove the linker and coiled-coil domain, having only the SH3 domain determining two CB isoforms (Fig. 4I-J). Analysis of CB Δ C_SH3+_ and CB Δ C_SH3-_ protein half-life revealed intermediate values between WT CB1_SH3-_ (1.8 hr) and WT CB2_SH3-_ (7.3 hr), around 4 hr. This result suggests that the C-terminus sequence and possibly the tertiary protein structure to be major determinants of CB protein stability.

### CB sequesters Cdc42 in the absence of gephyrin

Although CB has been described as a GABAergic synapse-specific RhoGEF with Cdc42 as preferred substrate, their functional relationship within neurons remains unclear. As Cdc42 influence on cytoskeleton organization is well documented [25], we wondered whether CB modulation of Cdc42 could somehow impinge on actin reorganization. To test this possibility, we co-transfected eGFP-CB2 Δ PH, eGFP-CB1_SH3-_ or eGFP-CB2_SH3-_ along with VSVG-Cdc42 in HEK-293T cells and examined actin reorganization 12 hr post-transfection using morphological analysis. We stained actin using phalloidin and counted filopodia in co-transfected HEK-293T cells (Fig. 5A-C). Comparison of cells transfected with eGFP, eGFP-CB1_SH3+_, eGFP-CB1_SH3-_, eGFP-CB2_SH3+_ or eGFP-CB2_SH3-_ revealed an increase in filopodia formation when eGFP-CB isoforms expressed (Fig. 5D; One-way ANOVA, P< 0.0001, Tukey multiple comparison test). This observation was unexpected, given that Cdc42CA causes membrane ruffling and Cdc42DN leads to filopodia formation (Fig. 5E-G).

**Figure 5:**
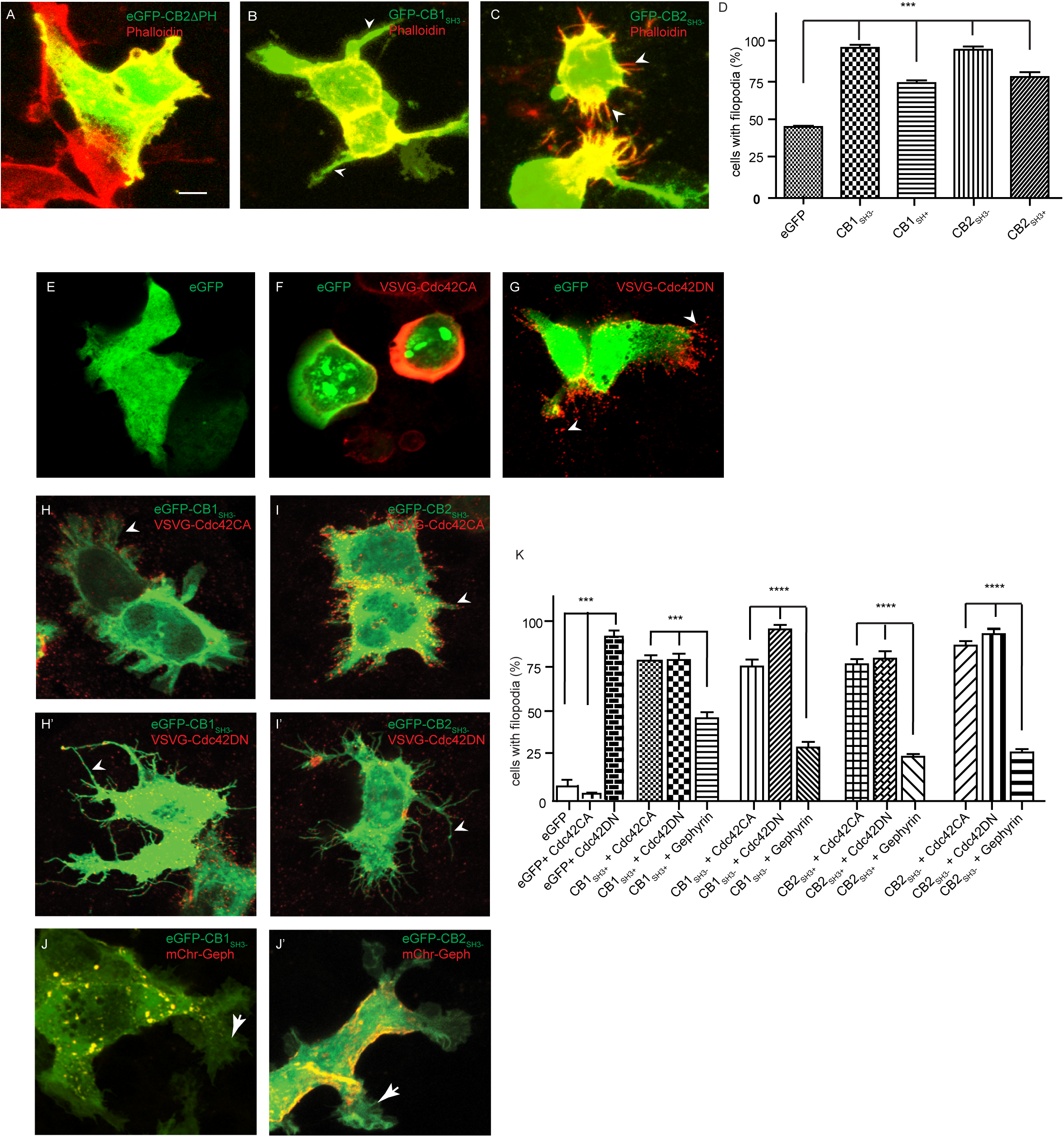
CB1 and CB2 sequester Cdc42 and influence actin dynamics. (A-C) Morphology of HEK-293T cells transfected with eGFP-CB2ΔPH, eGFP-CB1_SH3-_ or eGFP-CB2_SH3-_ and stained for actin using phalloidin. **(D)** Quantification of filopodia in HEK-293T cells expressing eGFP, eGFP-CB1_SH3+_, eGFP-CB1_SH3-_, eGFP-CB2_SH3+_ or eGFP-CB2_SH3-_ shows an increase in cells expressing CB isoforms. **(E-G)** HEK-293T cell morphology after transfection with eGFP, eGFP and VSVG-Cdc42 (CA) or VSVG-Cdc42 (DN). Arrow heads show accumulation of VSVG-Cdc42 (DN) (red) in filopodia. **(H-I)** HEK-293T cells co-transfected with VSVG-Cdc42 (CA) and eGFP-CB1_SH3-_ or eGFP-CB2_SH3-_. Arrow heads show accumulation of VSVG-Cdc42 (CA) (red) in filopodia. **(H’-I’)** HEK-293T cell morphology after co-transfection with VSVG-Cdc42 (DN) and eGFP-CB1_SH3-_ or eGFP-CB2_SH3-_ showing enhanced filopodia formation. Arrow head show accumulation of VSVG-Cdc42 (DN) (red) in filopodia. **(J-J’)** HEK-293T cell morphology after co-transfection with mCherry-gephyrin and eGFP-CB1_SH3-_ or eGFP-CB2_SH3-_. Arrow show membrane ruffling instead of filopodia. **(K)** Quantification of filopodia in HEK-293T cells upon co-expression of VSVG-Cdc42CA, VSVG-Cdc42DN or mCherry-gephyrin along with V5-CB isoforms.

In order to clarify how CB might induce filopodia in HEK-293T cells, we co-transfected either VSVG-Cdc42CA together with eGFP-CB1_SH3-_ or eGFP-CB2_SH3-_. Co-transfection of eGFP-CB isoforms suppressed VSVG-Cdc42-CA phenotype and induced filopodia formation (Fig. 5H-I), suggesting that CB might sequester Cdc42 irrespective of its activation status and prevent its interaction with actin. We confirmed this hypothesis by co-transfecting a dominant negative (DN) VSVG-Cdc42 mutant along with eGFP-CB1_SH3-_ or eGFP-CB2_SH3-_ isoforms (Fig. 5 H’-I’). While VSVG-Cdc42-DN promoted filopodia formation upon transfection, co-expression of eGFP-CB isoforms amplified this phenotype, suggesting a causal link between CB-mediated inactivation of Cdc42, likely owing to Cdc42 sequestration. We have reported earlier that CB, Cdc42, and gephyrin can form a ternary protein complex [15]. Hence, we next tested whether gephyrin co-transfection might influence eGFP-CB1_SH3-_ or eGFP-CB2_SH3-_-mediated Cdc42 sequestration. Interestingly, and in line with a role for CB regulating Cdc42 at postsynaptic sites, mCherry-gephyrin co-expression reduced filopodia formation when the V5-CB1_SH3-_ or V5-CB2_SH3-_ isoforms were expressed. Instead, the HEK-293T cells showed ruffled membrane, indicative of Cdc42 activity (Fig. 5J-J’). Quantification confirmed that gephyrin co-expression along with VSVG-CB isoforms significantly reduces filopodia formation (Fig. 5K, One-way ANOVA, P<0.001; Tukey multiple comparison test).

To explore further the relationship between CB and Cdc42 we turned to biochemistry. In this assay, we purified bacterially overexpressed GST-Cdc42 and incubated it with γsGTP or GDP to mimic active and inactive forms, respectively. Next, we incubated lysate of HEK-293T cells overexpressing V5-CB isoforms with immobilized GSTCdc42 γsGTP or GDP. Pull down of GST-Cdc42, followed by WB for V5-CB, showed CB interaction with both active and inactive forms of Cdc42 (Suppl. Fig. 2A-B). This result confirms that CB can interact with both active and inactive forms of Cdc42.

If gephyrin interaction determines CB activation of Cdc42, then we should find a reduced Cdc42 interaction with CB in gephyrin co-expressing cells. To test this possibility, we co-transfected VSVG-Cdc42, V5-CB1_SH3-_ or V5-CB2_SH3-_ with or without FLAG-gephyrin (Suppl. Fig. 2C). We bacterially expressed and purified GST-Pak1 binding domain (GST-PKB). GST-PKB interacts with active Cdc42 with high specificity and affinity [26]. Hence, we depleted free activated Cdc42 from HEK-293T lysate by incubating with GST-PKB immobilized on glutathione agarose beads (Suppl. Fig. 2D, bottom panel). We collected active-VSVG-Cdc42 depleted HEK-293T cell supernatant and immunoprecipitated VSVG-Cdc42 using an antibody against VSVG. WB for V5 allowed us to determine the relative levels of Cdc42 bound to V5-CB1_SH3-_ or V5-CB2_SH3-_ in the presence or absence of gephyrin (Suppl. Fig. 2C). These experiments confirmed the reduced Cdc42 interaction with V5-CB1_SH3-_ and V5-CB2_SH3-_ when FLAG-gephyrin was co-expressed. Taken together, these results indicate that, rather than activating Cdc42, CB isoforms prevent its activation (by other GEFs) by sequestrating it. In turn, gephyrin binding the CB reduces the amount of sequestered Cdc42 and thereby the availability of Cdc42 for remodeling of the actin cytoskeleton.

### CB2 influences neuronal maturation by Cdc42 sequestration

Biochemical experiments have shown consistently a stronger interaction between Cdc42 and CB2 than CB1, both *in vitro* and in transfected HEK-293T cells. Therefore, the impairment of dendrite growth observed in adult newborn neurons upon eGFP-CB2_SH3-_ overexpression could be due to Cdc42 sequestration. To confirm this idea, we infected newborn neurons with retroviruses expressing eGFP-Cdc42DN and quantified migration distance of these neurons and dendrite formation at 14, 28 and 42 dpi (Fig. 6A-A”). We did not observe any significant differences in the migration of cells expressing eGFP or eGFP-Cdc42DN, in line with the results presented in Figure 3. However, we observed a significant reduction in dendrite complexity at 28 dpi in newborn neurons expressing eGFP-Cdc42DN (Fig. 6B-B”; Mann Whitney test for the AUC P = 0.0226 for B’). This phenotype of eGFP-Cdc42DN overexpression was similar to that produced by eGFPCB2_SH3-_ overexpression, albeit less severe.

**Figure 6:**
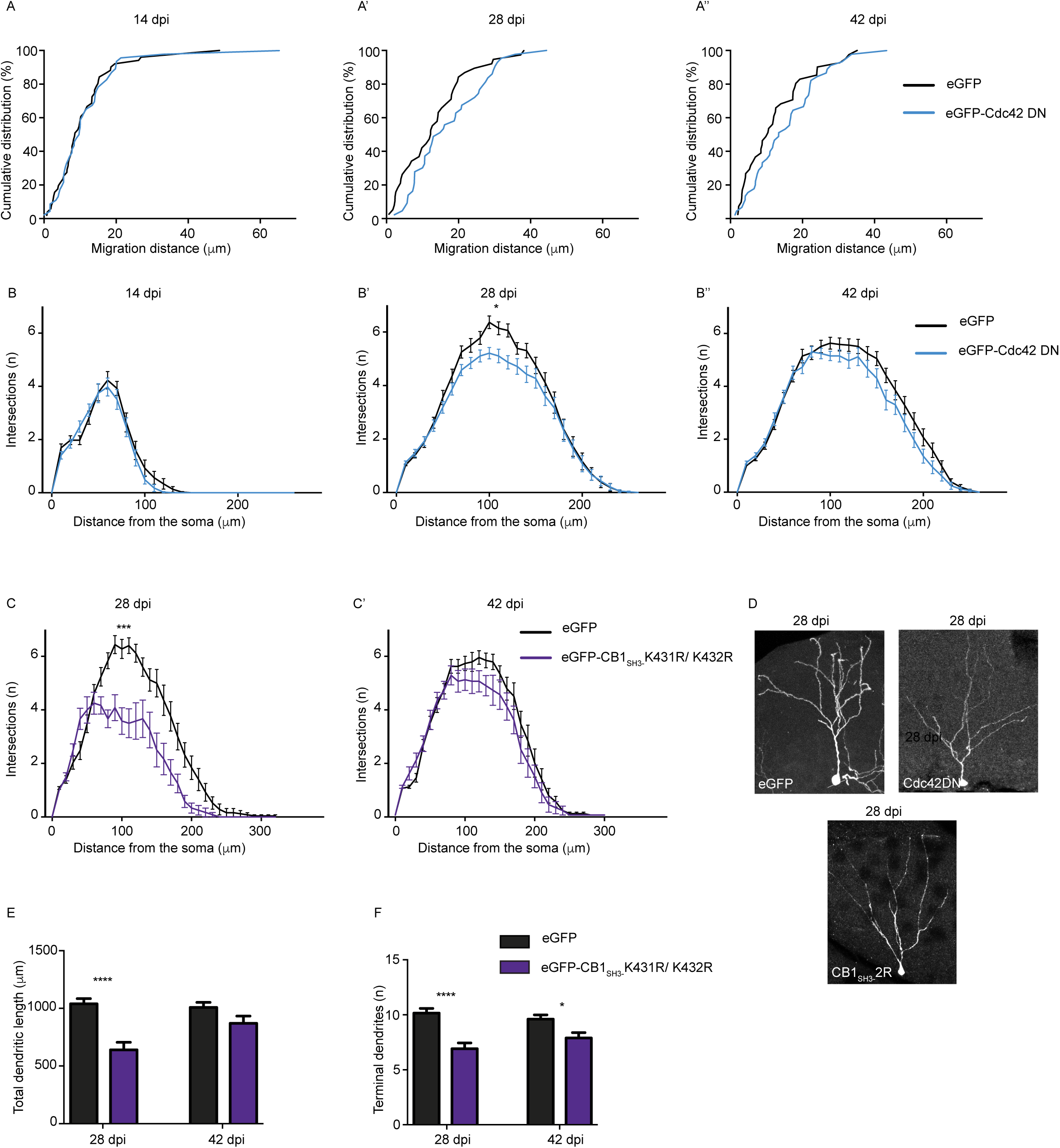
Cdc42DN and CB1 Ub mutant behave like CB2_SH3-_ *in vivo*. (A-A”) Adult newborn neurons expressing eGFP-Cdc42 (DN) do not show any migration defects at 14 dpi, 28 dpi and 42 dpi. **(B-B”)** Adult newborn neurons expressing eGFP-Cdc42 (DN) show defect in dendrite maturation at 14 dpi, 28 dpi and 42 dpi. **(C-C’)** Adult newborn neurons expressing eGFP-CB1_SH3-_ 2R mutant exhibit reduced dendritic complexity at 28 dpi, but not 42 dpi. **(D)** Morphology of adult newborn neuron expressing eGFP, eGFPCdc42 (DN) or eGFP-CB1_SH3-_ 2R mutant. **(E)** Quantification of total dendrite length showing reduced length in neurons expressing eGFP-CB1_SH3-_ 2R mutant at 28 dpi, but not 42 dpi. **(F)** The number of terminal dendrites is reduced in neurons expressing eGFP-CB1_SH3-_ 2R mutant at both 28 dpi and 42 dpi. Kolmogorov-Smirnov test (A-A”), N = 38-51 cells/group. Mann-Whitney t-test AUC (B-B”, C-C’), N = 12-41 cells/group. Two-way ANOVA, Bonferroni post-hoc test (E,F), N = 12-25 cells/group.

In view of this finding, we wondered whether the shorter protein half-life for eGFP-CB1_SH3-_ restricted its function when overexpressed in newborn neurons. If this were true, then expressing the eGFP-CB1_SH3-_ K431R/432R mutant, which exhibits a half-life similar to eGFP-CB2_SH3-_, should also impair dendritic aborization. Hence, we infected newborn neurons with eGFP or eGFP-CB1_SH3-_ K431R/432R mutant retrovirus, and looked for defects in dendritic arborization. Sholl analyses showed reduced dendritic complexity in cells overexpressing eGFP-CB1_SH3-_ K431R/432R at 28 dpi (Fig. 6C-C’, Mann Whitney test for AUC, P = 0.0004 for B’). Furthermore, we also observed a significant reduction in the total dendritic length and a reduction in the number of terminal branches in these neurons (Fig. 6D-F). These observations confirm that protein half-life is a major determinant of CB1 and CB2 isoform functional differences.

Given that CB1 differentially influences gephyrin clustering along the proximal-distal axis of dendrites in primary neurons (Fig. 1), we wondered whether this effect also was influenced by CB1 stability. We co-transfected DIV 8 neurons with eGFP-gephyrin and V5-CB1_SH3+_, or V5-CB1_SH3-_, or V5-CB1_SH3+_ 2R mutant, or V5-CB1_SH3-_ 2R mutant, and analyzed the density of postsynaptic eGFP-gephyrin clusters along the proximal-distal axis at DIV8 + 7 (Suppl. Fig. 3A-E’). Neurons co-expressing either V5-CB1_SH3+_ or V5-CB1_SH3-_ showed significantly increased eGFP-gephyrin clustering at distal dendritic segments. However, in neurons co-expressing V5-CB1_SH3+_ K491R/492R or eGFP-CB1_SH3-_ K431R/432R we saw an overall increase in gephyrin clustering without any specific gradient along the proximal-distal axis (Suppl. Fig. 3F-H). Therefore, these data identify a further role for protein stability in the functional differentiation between CB1 and CB2 isoforms.

### CB knockdown mimics CB1 and CB2 defect in newborn neurons

Upon overexpression in adult-born neurons, our data identifies a role for CB1 in neuronal cell migration and for CB2 in dendrite growth. In order to rule out overexpression artifacts, we reduced endogenous CB expression by PanCB shRNA-mediated silencing to unmask its contribution for regulating neuronal migration and dendrite maturation. We tested several PanCB shRNA sequences in HEK-293T cells and incorporated the most effective sequence in a retrovirus IRES GFP to label infected newborn neurons. We analyzed for migration differences at 21 dpi and 42 dpi as our earlier experiments showed strongest effects during these time points. As control, eGFP alone or scrambled shRNA sequence was used. Expression of PanCB shRNA reproduced the CB1_SH3-_ overexpression phenotype on neuronal migration. Fewer than 70% PanCB shRNA-expressing cells had migrated up to 15 μm at 21 dpi whereas cells expressing either eGFP or scrambled shRNA control had migrated more than 25 μm from the border of the SGZ (Fig. 7A-A’). This finding confirms the importance of a delicate regulatory balance to maintain correct levels of activated Cdc42.

**Figure 7:**
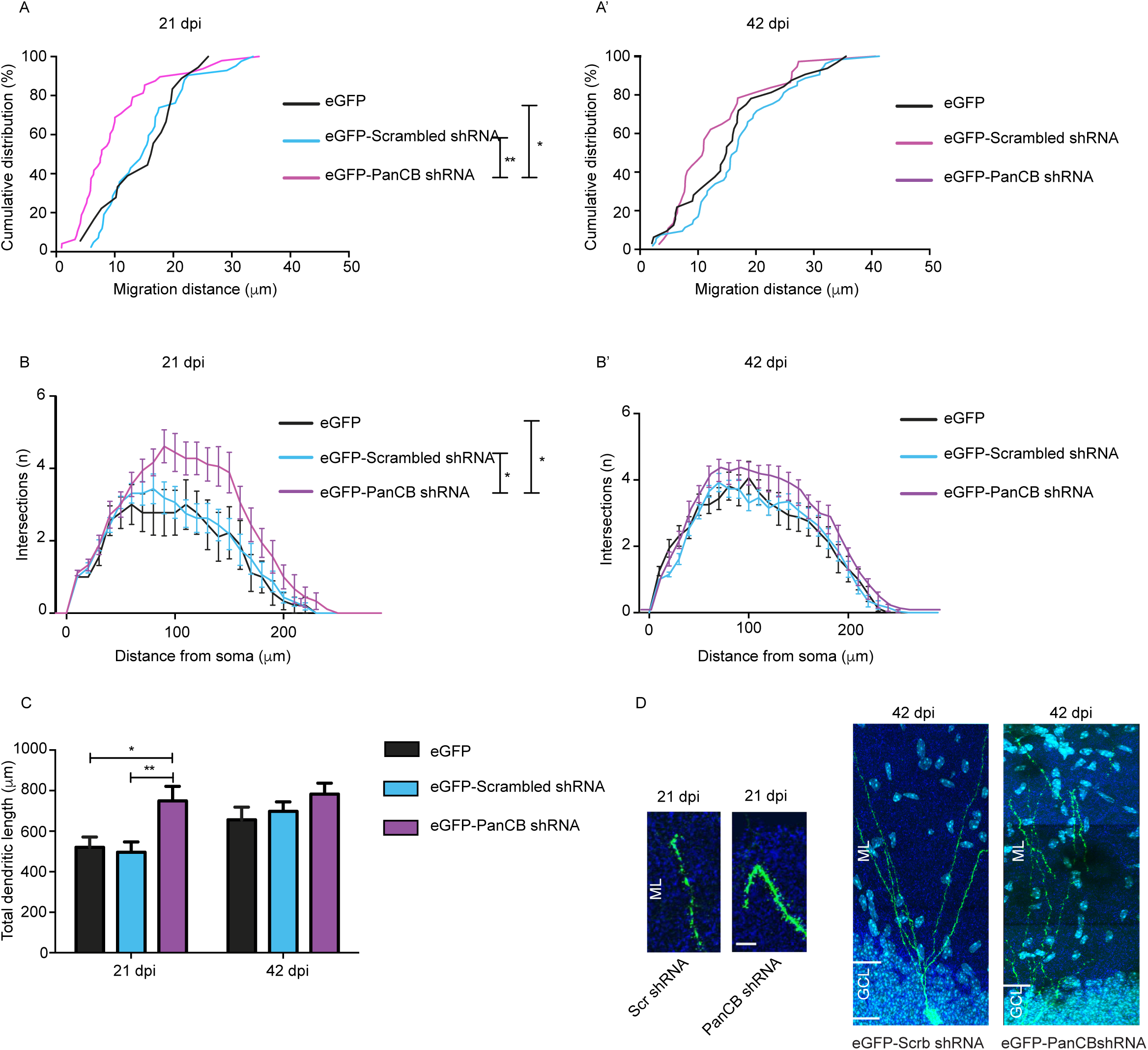
PanCB shRNA expression in adult newborn neurons recapitulates CB1 and CB2 functions. (A-A’) Adult newborn neurons expressing eGFP-PanCB shRNA migrate less at 21 dpi compared to eGFP and eGFP-scrambled shRNA expressing cells. There is no migration difference between eGFP, eGFP-scrambled shRNA or eGFPPanCB shRNA expressing cells at 42 dpi. **(B-B’)** Adult newborn neurons expressing eGFP-PanCB shRNA show reduced dendritic complexity at 21 dpi in comparison to eGFP or eGFP-Scrambled shRNA controls. There is not maturation difference between eGFP, eGFP-Scrambled shRNA or eGFP-PanCB shRNA at 42 dpi. **(C)** Total dendrite length is increased in neurons expressing eGFP-PanCB shRNA compared to eGFP or eGFP-Scrambled shRNA. **(D)** Morphology of adult newborn neurons expressing eGFP-Scrambled shRNA or eGFP-PanCB shRNA at 21 dpi and 42 dpi. GCL, granule cell layer; ML, molecular layer. Scale Bar 10 μm. One-way ANOVA, Bonferroni post-hoc test (A-A’), N = 18-53 cells/group. One-way ANOVA, Bonferroni post-hoc test AUC (BB’), N = 9-26 cells/group. Two-way ANOVA, Bonferroni post-hoc test (C), N = 12-23 cells/group.

Interestingly, PanCB shRNA-expressing neurons showed increased dendritic complexity at 21 dpi, which is opposite of the eGFP-CB2_SH3-_, eGFP-Cdc42DN or eGFP-CB1_SH3-_ K431R/432R mutant overexpression phenotypes (Fig. 7B-B’; One-way ANOVA for AUC, P = 0.0154 for B). We confirmed this finding by analyzing the total dendritic length in GCs expressing PanCB shRNA (Fig. 7C). We even observed distal branches in PanCB shRNA positive cells to revert their radial course when reaching the outer border of the ML, in both the upper and lower blades of the DG (Fig. 7D). These results demonstrate that overexpression of CB1 or CB2 isoform into newborn neurons limit dendrite growth presumably via sequestration of Cdc42.

### CB1_SH3-_ can rescue gephyrin scaffolding defect in adult-born neurons of *Gabra2* KO mice

We have reported that CA1 pyramidal cells in *Gabra2* KO mice exhibit a reduced frequency of miniature IPSCs, but no change in GABAergic current amplitude [27]. Hence, we infected adult-born neurons in *Gabra2* KO mice using a eGFP retrovirus and analyzed for gephyrin clustering defects along the proximal-distal axis at 14, 28 and 42 dpi (Fig. 8A-B’). We examined α1 subunit expression to determine whether this GABA_A_R subtype compensates in *Gabra2* KO cells. Staining for the α1 subunit in eGFP-positive newborn neurons in *Gabra2* KO at 14, 28 and 42 dpi showed elevated α1 GABA_A_R levels at 28 dpi in both proximal and distal dendritic segments (Fig. 8C-C’). This increase of α1 subunit clusters was not seen at 42 dpi, suggesting a role for extrinsic factors in shaping inhibition during the maturation process.

**Figure 8:**
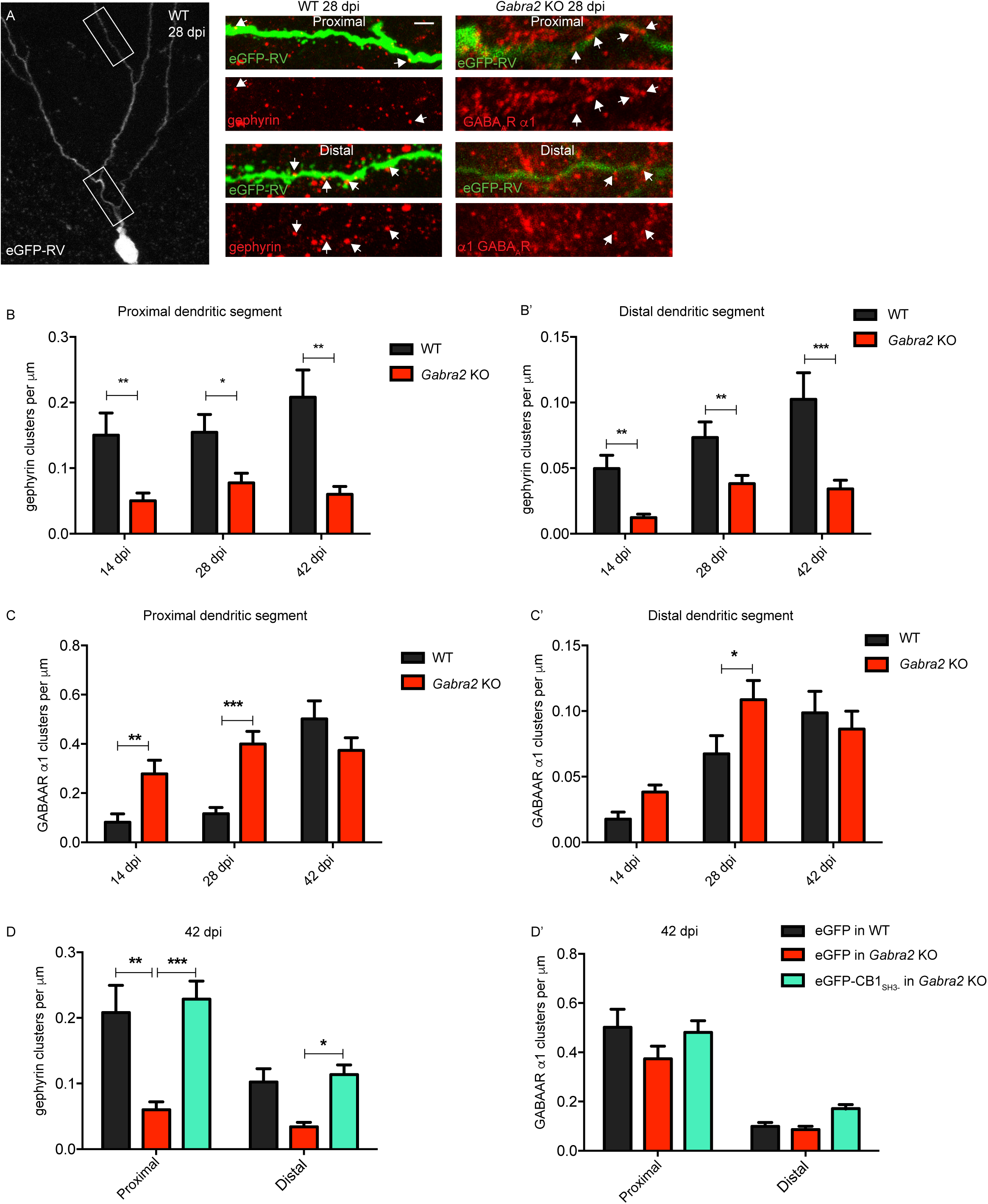
eGFP-CB1_SH3-_ can rescue gephyrin scaffolding in *Gabra2* KO neurons. (A) Morphology of retrovirus eGFP with proximal and distal dendritic sites. Right panels; WT and *Gabra2* KO neuron expressing eGFP with gephyrin and α1 GABA_A_R staining. Arrows show reduced gephyrin clustering and increased α1 GABA_A_R clusters in *Gabra2* KO neurons. **(B-B’)** α1 GABA_A_R cluster density in proximal dendritic segments show significant increase in *Gabra2* KO cells at 14 dpi, 28 dpi, but not 42 dpi. Similarly, α1 GABA_A_R cluster density in distal dendritic segments are increased *Gabra2* KO cells at 28 dpi, but not 14 dpi and 42 dpi. **(C-C’)** Gephyrin cluster density in proximal and distal dendritic segments show significant reduction in *Gabra2* KO cells at 14 dpi, 28 dpi and 42 dpi. **(D-D’)** Gephyrin clustering at both proximal and distal dendritic segments is restored in *Gabra2* KO neurons upon eGFP-CB1_SH3-_ overexpression. Rescue of gephyrin clustering does not influence α1 GABA_A_R cluster density in *Gabra2* KO neurons.

α2 GABA_A_R can directly interact with CB and facilitate gephyrin scaffold recruitment at GABAergic postsynaptic sites [12,28]. Hence, we examined in *Gabra2* KO mice, where we have reported deficit in gephyrin scaffolding both in hippocampal formation and neocortex [20,27], whether CB overexpression can compensate for the absence of the GABA_A_R in facilitating gephyrin clustering. We quantified gephyrin cluster density in the first 40 μm segment of dendrite from the soma and found a significant reduction in GCS from *Gabra2* KO compared to the WT mice. Similar observations were made in distal dendritic segments, as well (80-120 μm) (Fig. 8B-B’).

To test whether CB1_SH3-_ overexpression could rescue endogenous gephyrin scaffolding in *Gabra2* KO cells, we overexpressed eGFP-CB1_SH3-_ using a retrovirus in *Gabra2* KO newborn neurons and analyzed for gephyrin clustering in the proximal and distal dendritic segments at 42 dpi. eGFP-CB1_SH3-_ overexpression consistently rescued endogenous gephyrin clustering in GCs from *Gabra2* KO mice (Fig. 8D-D’).

### CB1_SH3-_ rescues migration defects in adult newborn neurons from *Gabra2* KO mice

We have reported a direct link between the *Gabra2* gene and adult neurogenesis [6]. Hence, we wanted to examine whether CB1_SH3-_ function occurs downstream of α2 GABA_A_R and infected WT and *Gabra2* KO mice with eGFP retrovirus to analyze cell migration differences at 14, 28 and 42 dpi. α2 GABA_A_Rs contribute towards cell migration, as in the absence of the *Gabra2* gene product, newborn neurons migrate 60% longer distance away from the SGZ. This difference was highly significant at 14 dpi (Suppl. Fig. 4A-B”; Kolmogorov-smirnov test, p<0.0001), but diminished over time, as the WT cells caught up at 28 and 42 dpi.

If CB1_SH3-_ functions downstream of α2 GABA_A_Rs, we reasoned that it should be able to reverse the migration phenotype observed in adult-born neurons of *Gabra2* KO mice. We infected *Gabra2* KO mice with either eGFP or eGFP-CB1_SH3-_ retrovirus and quantified migration of labeled GCs at 14, 28 and 42 dpi. The expression of eGFP-CB1_SH3-_ reduced the migration of the newborn neurons by 60% shorter distance from the SGZ. The migration difference did not normalize at later time points (Suppl. Fig. 4C-C”; Kolmogorov-Smirnov test, P = 0.0044, 0.0274, 0.0096 respectively). This confirms a role for CB1_SH3-_ downstream of α2 GABA_A_Rs and highlights the relevance of CB-mediated regulation of Cdc42 function for proper cell migration.

## Discussion

The results of this study reveal distinct functional differences between CB1 and CB2 splice isoforms that can be abolished by mutating two lysine residues in the C-terminus of CB1 and thereby prolonging the half-life of CB1 by preventing ubiquitination and subsequent degradation. In primary neurons, CB1 and CB2 differentially promote the formation of gephyrin clusters (hence, GABAergic synapses) depending on the degree of maturity of dendritic segments. Further, the data show that CB isoforms indirectly impinge upon the actin cytoskeleton, in addition to their role at GABAergic synapses, to regulate migration of immature adult-born neurons in the dentate gyrus and the development of their dendritic tree. The multiple functions of CB isoforms are likely mediated by Cdc42, but, contrary to expectancy, CB isoforms appear to be negative regulators of Cdc42 activity, and act by trapping Cdc42 in a gephyrin-dependent manner. In line with this finding, no effect of adding/removing the SH3 cassette was observed on the CB functions investigated in this study. As the SH3 domain regulates the catalytic activity of CB, the latter is not directly involved in these functions of CB. Silencing CB expression in adult-born dentate gyrus GCs reveals that endogenous CB promotes neuronal migration away from the SGZ and limits dendritic growth in immature GCs. Finally, CB overexpression in adult-born GCs from *Gabra2* KO mice reverts deficits in migration and gephyrin postsynaptic clustering observed in the absence of α2-GABA_A_Rs, suggesting that CB acts downstream of GABA_A_Rs, even for its non-synaptic functions.

### Developmental regulation of CB mRNA splicing

Our RTqPCR analysis revealed differential regulation of CB splicing during development, favoring CB1 isoform expression at early stages of development, especially in the neocortex. Assuming equal mRNA translation, these data favor a role of CB1 in immature neurons, possibly at the onset of synaptogenesis. However, the short half-life of CB1 relative to CB2 suggests a transient role for this isoform. The half-life of CB1 determines its specific role even upon overexpression in primary neurons, as seen upon mutation of C-terminal lysine residues that render CB1 CB2-like. Therefore, we conclude that ubiquitination of CB1 is a novel mechanism restricting its functional repertoire. It would be interesting to test whether it regulates the interaction of CB1 with Ccd42, which is overall weaker than that of CB2 with Cdc42.

A pyramidal cell-specific transcriptional program involving transcription factor NPAS4 has been reported to coordinate the distribution of GABAergic synapses by increasing the number synapses on the cell body, while decreasing the synapse number on the apical dendrite [29]. In addition, neuronal cell-specific expression of splicing factors SLM1 and SLM2 has been shown to contribute towards the postsynaptic biochemical diversity [30]. Hence, identification of CB1-facilitated gephyrin clustering selectively in the distal portion of immature dendrites adds to the repertoire of biochemical diversity at GABAergic synapses, complementing the compartment-specific interneuron connectivity.

### Ubiquitination as determinant of CB isoform function

The role of the ubiquitin proteasome system (UPS) in synaptic plasticity is an emerging concept at inhibitory synapses. However, at glutamatergic synapse protein ubiquitination and degradation are important determinants of synapse function [31]. In the current study, we highlight the importance of CB1 protein turnover for compartment-specific eGFP-gephyrin clustering. This finding implicates the local availability of CB1 protein pools as a basis for GABAergic synapse distribution along the elongating neuronal dendrite. We also report the importance of UPS in adult newborn neurons *in vivo* to explain the GABAergic basis for actin regulation and dendrite outgrowth. However, the distribution and regulation of UPS in developing neurons are currently not understood and require further investigations.

Strikingly, in wildtype mice, CB2_SH3-_ overexpression and shRNA-mediated downregulation of all CB isoforms had opposite effects on dendrite arborization, suggesting that CB2 activates signaling factors to regulate dendritic growth. A role for small Rho-GTPase, such as Cdc42 (or possibly TC10) downstream of CB2 might explain the enhanced dendrite outgrowth phenotype upon CB silencing. This hypothesis implies, however, that CB inhibits Cdc42 activation. Our data confirms this hypothesis by demonstrating the failure of dendrites to stop growing when they reach the outer surface of the molecular layer upon CB down regulation (Fig. 7D).

### GABA_A_R-mediated regulation of adult neurogenesis

Evidence supports the notion that specific GABA_A_R subtypes, determined by their α subunit variants, regulate distinct stages of adult neurogenesis in the dentate gyrus [6, 32, 33]. Thus, strong genetic and pharmacological evidence points towards the regulation of cell-fate determination by α5 GABA_A_Rs, whereas proliferation on precursor cells is modulated by α4 GABA_A_Rs and migration of adult-born GCs away from the SGZ is regulated in opposite directions by α4-and α2 GABA_A_Rs [6]. Finally, dendritic growth and arborization, as well as GABAergic synapse formation, are modulated, at least in part by α2 and α5 GABA_A_Rs [6,33]. Our present results implicate CB1 and CB2 downstream of α2 GABA_A_Rs for both GC migration and postsynaptic gephyrin clustering, providing a mechanistic explanation how GABAergic synaptic transmission might act on the actin cytoskeleton regulating neuronal mobility and dendrite formation.

### CB negatively regulates Cdc42 activation

Overexpression experiments can be challenging to interpret, because the effects might be minimized or even reverted due to exhaustion of an essential factor present at a limited level. In this context, the paradoxical observation that the inhibitory effect of CB1 on migration of adult-born GCs could be mimicked upon overexpression of a DN Cdc42 mutant provided the first clue towards the possibility that CB overexpression might affect the regulation of Cdc42 activation. More direct evidence for an inhibitory effect of CB isoforms on actin remodeling mediated by Cdc42 was obtained, therefore, in HEK-293T cells, where the formation of filopodia and membrane ruffling are robust and well-established readout of Cdc42 inactivation and activation, respectively [34]. The results of these experiments are best explained by assuming that binding of Cdc42 to CB limits (or prevents) its activation, presumably by other GEFs like TC-10 present in HEK-293T cells. Importantly, this interaction between CB and Cdc42 is modulated by gephyrin, providing an elegant mechanism to explain how CB action on the cytoskeleton at postsynaptic sites might be locally restricted.

Our pull-down experiments confirmed that CB2 interacts stronger than CB1 with Cdc42 (possibly due to CB1 ubiquitination, see above) and that CB1 and CB2 bind to both activated (i.e., GTP-bound) and inactive Cdc42 (GDP-bound). These biochemical observations are compatible with the hypothesis that, albeit being a weak activator of Cdc42, CB mainly acts upon the actin cytoskeleton by limiting its availability for other activators of this ubiquitous small GTPase.

## Conclusions

Taken together, our results provide a new framework for the differential functions of CB isoforms in neurons. CB1 is an early expressed, short-lived isoform regulated by ubiquitination. It modulates GABAergic synapse formation and regulates neuronal migration downstream of α2 GABA_A_R. The latter effect involves downregulation of Cdc42 activity by trapping it away from other Cdc42 activators. CB2 is a constitutively expressed variant, with a longer half-life, which promotes dendrite growth and GABAergic synapse formation. The action of CB2 also involves negative regulation of Cdc42 activation, but, based on its distribution in adult CNS [12], might be localized mainly at GABAergic postsynaptic sites. The Ubiquitin-Proteasome System (UPS), by limiting the action of CB1 (and possibly its binding to Cdc42), thus appears to be a major regulator of the function of the GABAergic system.

## Materials and Methods

### Plasmids

CB isoforms CB1_SH3+_, CB1_SH3-_, CB2_SH3+_ and CB2_SH3-_ were cloned from rat whole brain RNA using primers Fwd: 5’-ATGCAGTGGATTAGAGGC-3’; Rev: 5’-CTAATAGTGCCATTTTCTTTGG-3’ for CB1 isoforms, and Fwd: 5’-ATGCAGTGGATTAGAGGC-3’; Rev: 5’-CGCTAAGCTTCATGACTCTGCT GATCA-3’ for CB2 isoforms. mCherryC2-CB1SH3 +, mCherryC2-CB1_SH3-_, mCherryC2-CB2_SH3+_, mCherryC2-CB2_SH3-_, expression vectors were generated by subcloning CB into eGFPC2 vector backbone using HindIII and BamHI. eGFP sequence was later replaced by mCherry sequence using NheI and XhoI sites. pCR3-V5-CB1_SH3+_, pCR3-V5-CB1_SH3-_, pCR3-V5-CB2_SH3+_ and pCR3-V5-CB2_SH3-_ were generated by subcloning the V5 tag into the pCR3-CB vectors using HindIII restriction site. The eGFP-gephyrin P1 variant has been described previously (Lardi-Studler et al 2007). FLAG-gephyrin is described previously (Tyagarajan et al 2011). pGEX-2T-GST Cdc42WT was obtained from Addgene (#12969) and pGEXTK-Pak1 (70-117) was obtained from Addgene (#12117). pRKVSV-Cdc42 WT, pRKVSV-Cdc42 DN (N17), pRKVSV-Cdc42 CA (QL) were gifts from Prof. Kenneth Yamada (NIH). CB mutants were generated using site directed mutagenesis according to vendor manual (Agilent Technologies, USA) using pCR3-V5-CB splice isoforms as the template. The truncated V5-CBΔC_SH3+_ and V5-CB Δ C_SH3-_ were generated by inserting a Stop codon after Ile 440 (Ile380 for CB_SH3-_, respectively) into pCR3-V5-CB1 isoforms. HA-ubiquitin was a gift from Dr. Teier, Heidelberg, Germany.

### Primary neuron culture

Primary hippocampal neuron cultures were prepared as described previously in [35]. Hippocampal cultures were transfected with eGFP-gephyrin and mCherry-CB or V5-CB construct (500 ng of each plasmid), using a combination of Lipofectamine 2000 (Life Technologies) and CombiMag (OZ Biosciences). The neurons were grown in 2 mL of growth media for 8 days prior to transfection. The transfection mix was incubated at room temperature for 15 min before adding to the neurons. The transfection was stopped 25 min later by transferring the coverslips into a fresh 12 well dish containing the conditioned media.

### HEK-293T cell culture

HEK-293T cells were cultured at 37°C under a 5% CO_2_ atmosphere in Dulbecco’s modified Eagle’s medium (DMEM) supplemented with 10% fetal bovine serum (FBS). They were transfected with 2-3 μg DNA at 14-16 hr post-plating using polyethylenimine (PEI) according to the manufacturer’s recommendation. The whole cell lysate was prepared 24 hr post-transfection using EBC buffer containing Complete-mini (Roche) and a phosphatase inhibitor cocktail (Sigma).

### Animals

All experiments were performed in accordance with the European Community Council Directives of November 24, 1986 (86/609/EEC) and approved by the cantonal veterinary office of Zurich. Wild type C57Bl6/J-Crl1 mice were purchased from Charles River Laboratories (Germany). *Gabra2* KO mice were bred at the Institute of Pharmacology and Toxicology, University of Zurich and genotyped as described in [27].

### Retrovirus production

For retrovirus production, a non-replicative vector was adapted from the Moloney murine leukemia virus (MMLV). DNA constructs with the gene of interest (GFPCB1_SH3-_, eGFP-CB2_SH3-_, eGFP-CB1_SH3-_ 2R were cloned into the pCAG-V-PRE-eGFP vector. The eGFP-Cdc42DN and pCMV-gp expressing gag/pol genes for virus packaging and pCMV-vsv-g used as envelope protein plasmids were a kind gift from Prof. Sebastian Jessberger (University of Zurich).

ShRNA against ArhGEF9 were ordered from Origene Technologies (Rockville, USA) in a HuSH pRFP-C-RS (TF517554) backbone. GFP-panCB shRNA CTGATGAAGGACAGCCGCTATCAACACTT and 29-mer scrambled ShRNA cassette in pRFP-C-RS (TR30015) was used as control. pCAG-V-PRE-eGFP-panCB shRNA was subcloned using PstI restriction sites introduced using PCR primer 5’ to U6 promoter and 3’ to termination codon.

HEK-293T cells were transfected with the “CalPhos Kit” viral transfection protocol with three separate plasmids containing the capsid (CMV-vsvg), viral proteins (CMV @ gag/pol) and the gene of interest. Medium containing the virus was collected and the virus was purified with ultracentrifugation, re-suspended in cold phosphate-buffered saline (PBS) and stored at-80°C. The virus titer of minimum 10^6^ cfu/mL was used for sterotactic injections, after determination using serial dilution in HEK-293T cells.

The eGFP-CB1_SH3-_ K431/K432R (2R) viral vector and viral vector plasmid were generated at the Viral Vector Facility (VVF) of the Neuroscience Center Zurich.

### Stereotaxic intrahippocampal injections

Mice 8-12 weeks old were anesthetized by inhalation of isoflurane (Baxter) in oxygen, injected intraperitoneally (i.p.) with 1 mg/kg buprenorphine (Temgesic, Essex Chemicals, Lucerne, Switzerland), and head-fixed on the stereotaxic frame (David Kopf Instruments). The Nanoject II^TM^ Auto-Nanoliter Injector (Drummond Scientific Company) was used to deliver 69 nL of MMLV bilaterally into the hilus of the dentate gyrus 15 x (antero-posterior =-2 mm; lateral = ±1.5 mm; dorso-ventral =-2.3 mm, with the bregma as reference), under stereotaxic guidance (“Stereodrive software”). During the operation and recovery, the mice were held on a warm pad. After the operation they were received a second injection of buprenorphine.

### Tissue preparation for Immunohistochemistry

Mice were deeply anesthetized with i.p. injection of 50 mg/kg sodium pentobarbital (Nembutal) and perfused intracardially through the left ventricle with approximately 20-25 mL ice-cold, oxygenated Artificial Cerebrospinal Fluid (ACSF: 125 mM NaCl, 2.5 mM KCl, 2.5 mM CaCl2, 2 mM MgCl2, 26 mM NaHCO3, 1.25 mM NaH2PO4, 25 mM glucose; pH 7.4) at a flow rate of 10 mL/min [36]. The brain was immediately dissected and a block containing the hippocampal formation was fixed for 3 hr in ice-cold 4% paraformaldehyde solution (4% PFA in 0.15 M sodium phosphate buffer; pH 7.4). The tissue was rinsed with PBS and stored overnight at 4°C in solution of 30% sucrose in PBS for cryo-protection. The tissue was sectioned coronally after being frozen on tissue mounting fluid (M-1 Embedding matrix, Shandon, Thermo Scientific, USA), on the frozen (-40°C) block of a sliding microtome (MICROM HM 400, MICROM International GmbH, Walldorf, Germany). The tissue was cut into 40 μm thick sections, which were immediately transferred in ice-cold PBS as free-floating sections.

### Immunoprecipitation and western blot analysis

For immunoprecipitation 0.8 μl anti V5 antibody was added to 500 μl cell lysate and incubated over-night on a rotating wheel at 4°C and the protein complex was precipitated by using 50 μl of Protein A and Protein G Plus-Agarose beads (Calbiochem) in EBC buffer. The beads were washed once in EBC-based high-salt buffer (50 mM Tris-HCl pH 8.0, 500 mM NaCl and 1% Nonidet P-40 (Sigma-Aldrich) and twice in EBC buffer. After boiling the samples in 2xSDS sample buffer containing 15% 2-Mercaptoethanol (Bio-Rad) for 10 minutes at 72°C the supernatant was loaded onto SDS-polyacrylamide gels and run at 140V at room temperature. After transferring the protein bands onto PVDF membranes with constant 35 mA in Tris-glycine transfer buffer, WB were performed by blocking the membranes with 5% western blocking reagent (Roche Diagnostic) in Tris-buffered saline with Tween 20 (TBST) for 1 hr at room temperature and later incubating with primary antibody overnight at 4°C. Secondary antibody coupled to horseradish peroxidase or IRDye^®^ (LI-COR) was used to visualize the protein bands.

### Antibodies and immunohistochemistry

The following antibodies were used in this study: Chicken anti GFP (1:5000, Aves Labs Inc., USA), Mouse anti-V5 antibody (1:5000, Invitrogen, Carlsbad, USA) and (1:3000, Acris, SanDiego, USA), mouse anti-FLAG (1:5000, Sigma, Saint Louis, USA), mouse anti-actin (C4 clone) antibody (1:20000, EMD Millipore Corporation, Billerica, USA), rabbit anti-VGAT antibody (1:2000, Synaptic Systems, Göttingen, Germany), mouse anti-gephyrin antibody (mAb7a, 1:1000; or 3B11, 1:10000; Synaptic Systems, Göttingen, Germany), guinea pig anti-α1 subunit (home-made; [37], as well as a STrEP-tag Purification (IBA GmbH, Göttingen, Germany).

### Immunofluorescence of primary neuron cultures

Cells were fixed for 10 minutes in 4% PFA, rinsed in PBS and permeabilized with 0.1% Triton X-100 containing 10% normal goat serum. Immunohistochemistry was performed by incubating the cells with the primary antibodies diluted in PBS containing 10% normal goat serum for 60 min. After washing in PBS the cells were incubated with the secondary antibodies coupled to Cy3 or Cy5 (1:1000, Jackson ImmunoResearch) for 30 min. After drying the cells were mounted with fluorescent mounting medium (Dako Cytomation, Carpinteria, CA). Everything was performed at room temperature.

### Brain tissue staining

Sections were transferred into primary antibody solution (0.2% Triton X-100, 2% normal goat serum (NGS) and primary antibodies in PBS; pH 7.4) and incubated at 4°C, in darkness and under continuous agitation (100 rpm/min) for 72 hr. The sections were then washed 3x10 min in PBS and incubated in the secondary antibody solution (2% NGS and secondary antibodies targeting the species of the respective primary antibodies used in PBS; pH 7,4) at room temperature, in the darkness and under continuous agitation (100 rpm/min), for 6 h. DAPI 1:3000 was added into the secondary antibody solution to stain cell nuclei.

After incubation in the secondary antibody solution the sections were washed again 3x10 min in PBS, mounted onto gelatin-covered glass slides (Menzel, GmbH & Co KG), air-dried and coverslipped with Dako fluorescence mounting medium (Agilent Technologies, Santa Clara, CA, USA).

### RNA isolation and quantitative realtime-PCR

RNA was isolated from WISTAR rat brain sub regions as indicated at different ages (N = 5) using Sigma-Aldrich GenElute^TM^ Mammalian Total RNA Miniprep Kit; cDNA was prepared using random hexamers and SuperScript® II Reverse Transcriptase (Invitrogen). Quantitative PCR was performed on a 7900 HT Fast Real Time PCR system (Applied Biosystems). HOT FIREPol® EvaGreen® qPCR Mix Plus (ROX) (Solis Biodyne) was used with the designed primers to amplify cDNA (10ng/sample). Changes in mRNA levels were calculated using the ΔΔCt Method [38] relative to PanCB mRNA. Primer pairs common to all CB isoforms were designed and tested to normalize specific CB isoforms to PanCB levels. Bar charts and statistics were performed using Graphpad Prism. One-way ANOVA with Bonferroni post-hoc correction with significances: P<0.001 ***; P<0.01 **; P < 0.05 *

CB1 fwd 5’-GTAGGGTTGGAGAGGAAGAG-3’

CB1 rev 5’-TTGTGGTGGATAGGAAGGTG-3’

CB2 fwd 5’-GCACAACAAGGAAACCGAAGA-3’

CB2 rev 5’-TGGGTTACTTTCTGTTTAGACGCTTT-3’

panCB fwd 5’-CCCTGCTTCTTGGAGCATCA-3’

panCB rev 5’-TGATAGCGGCTGTCCTTCATC-3’

### Ubiquitination assay

HEK-293T cells were transfected with 1 μg of V5-CB and HA-ubiquitin using PEI as per vendor suggestions. Sixteen hours post-transfection, 5 μM MG132 (Tocris) was added and 5 hr post-treatment, cells were lysed for protein detection.

### Protein stability assay

V5-CB isoforms were transfected into HEK-293T cells using the PEI protocol. Sixteen hours after transfection, 100 μM cyclohexamide dissolved in culturing medium was added and the cells were lysed in EBC buffer (50 mM Tris-HCl pH 8.0, 120 mM NaCl and 0.5% Nonidet P-40) containing complete mini-protease inhibitor (Roche Diagnostics) and phosphatase inhibitor cocktail 2 and 3 (Sigma @ Aldrich) after 0, 1, 2, 4 or 8 hr of treatment for 20 min at 4°C. WB quantification of protein intensity was done using LI-COR odyssey scanner and image studio. The area under the curve was normalized to actin and a half-life curve was fitted using Graphpad Prism.

### Image analysis and quantification

For analysis of morphology of HEK-293T cells and hippocampal neurons as well as gephyrin and a1 subunit clustering in neurons, a confocal laser-scanning microscope (LSM700 or LSM710, Carl Zeiss AG, Jena, Germany) with a 25 x or 40 x oil immersion objective was used for image acquisition. The pinhole was set for all channels at 1 Airy unit; pixel size typically was 90 nm and a z-stack (3-4 steps at 0.5 μm interval) was acquired by sequential scanning of each channel. For each condition in *in vitro* experiments (primary neurons and HEK-293T cells), 15 cells per group from 3-4 independent experiments were imaged. For *in vivo* experiments, at least 3-5 mice per condition were used for analysis. In each mouse, images from 10-15 cells were acquired. Image analysis was performed using ImageJ (http://rsb.info.nih.gov/ij/).

### Cell migration analysis

In order to measure the migration distance, CSLM images were acquired with a 25 x oil immersion objective, in a way that the position of GFP-labeled granule cells could be clearly observed into the granular cell layer (GCL), labeled with DAPI. Using ImageJ, the orthogonal distance from the center of the cell to the base of the GCL was measured and compared using one-way ANOVA.

### Analysis of the dendritic tree

In order to image the neuronal cells on their whole, a 25 x oil immersion objective was used to acquire z-stack images in a slice interval of 1.5μm. The analysis was carried out using ImageJ, version 1.49o; Java 1.6.0_12 (Wayne Rusband, National Institutes of Health, USA). The *NeuronJ* plugin (NIH ImageJ, Meijering et al., 2004) was used to trace the dendritic trees and measure the length of the dendrites. In order to evaluate the complexity of the dendritic trees, Sholl analysis was performed [39]. The *Sholl Analysis* macro was used to count the number of intersections of the dendrites with imaginary concentric cycles, designed to have the middle of the cell soma as their center, and 10 μm subsequently increasing radii. The number of intersections for each given radius (and hence distance from the soma) were used to plot the diagrams, and the area under the curve (AUC) was used for statistical analysis.

### Analysis of postsynaptic eGFP-gephyrin clusters

For the analysis of synaptic clusters in primary neurons a 40x oil immersion objective (N.A. 1.4) was used and the pictures were acquired with a 1.8 zoom, a 0.45 μm inter-image interval and a 1024x pixels length in segments of proximal and distal dendrites. Images of proximal and distal dendritic segments of GFP-labeled cells were analysed with an ImageJ macro designed to identify eGFP-gephyrin clusters opposed to VGAT-positive terminals using threshold segmentation algorithms. The density of such postsynaptic clusters was normalized to a length of 20 μm.

Postsynaptic clusters in eGFP-labeled adult-born GCs were analyzed individually in each image of stacks covering the entire dendritic tree, using the same macro for identification of clusters by threshold segmentation.

### Statistical Analysis

Cumulative probability analysis was performed to analyze distribution of postsynaptic eGFP-gephyrin cluster size (Kolmogorov-Smirnov test) in primary neurons. The effect of CB isoforms on eGFP-gephyrin cluster density, CB isoform mRNA expression levels during brain development, CB effect on filopodia formation in HEK-293T cells and the effect of PanCB shRNA expression were analyzed by one way ANOVA with a Bonferroni post-hoc test. CB isoform effects on CB influence on dendritic arborization, the effect of Cdc42 sequestration by CB were analyzed using Mann Whitney test. All histograms, as well as the statistical analysis of the results of this study were carried out on Prism software (GraphPad Software Inc., La Jolla CA, USA).

## Acknowledgements

We thank late Abdul Mohammed Iqbal for his help with some of the biochemical assays.

## Figure Legends

**Fig. S1: CB1 and CB2 isoforms are Ub conjugated. (A)** HEK293T cells transfected with HA-Ubiquitin and V5-CB isoforms. The cells were treated with either DMSO or MG132 and analyzed for HA-Ub conjugation after IP for HA and WB for V5. Higher migrating V5 bands can be seen in MG132 treated samples. **(B)** HEK293T cells transfected with HA-Ub and V5-CB isoforms were treated with DMSO or MG132. IP for V5 followed by WB against HA-Ub showed enhanced Ub conjugated V5 CB isoforms in MG132 treated samples.

**Fig. S2: Biochemical analysis for CB and Cdc42 interaction. (A-B)** Bacterial overexpressed and purified GST-Cdc42 incubated with γsGTP or GDP to mimic active and inactive forms. GST, GST-Cdc42, GST-Cdc42 γsGTP or GST-CDC42 GDP incubated with HEK293T lysates overexpressing V5-CB1_SH3+_, V5-CB1_SH3-_, V5-CB2_SH3+_ or V5-CB2_SH3-_ showing interaction between active and inactive Cdc42 and CB isoforms. CB2 isoforms exhibit stronger interaction with Cdc42 compared to CB1 isoforms. **(A’-B’)** Protein loading controls showing equal amount of V5-CB isoform expression in HEK293T cells and GST-Cdc42 expression in bacteria. **(C)** HEK293T cells expressing vsvg-Cdc42 DN, vsvg-Cdc42 CA, vsvg-Cdc42 CA and V5-CB1_SH3-_ or V5-CB2_SH3-_ with or without FLAG-gephyrin. After pull down of free active Cdc42 from HEK293T cell lysate using GST-PKB we separated the supernatant for further analysis. We IP’ed vsvg-Cdc42 CA and performed WB against V5 to evaluate CB interaction with vsvg-Cdc42. Lanes containing FLAG-gephyrin co-expression showed reduced levels of CB interaction with vsvg-Cdc42 compared to CB alone lanes. **(D)** Pull down of free active vsvg-Cdc42 CA but not vsvg-Cdc42 DN using GST-PBD shows equal levels of active Cdc42CA in all lanes. Protein transfection levels in HEK293T cells are shown on the right panel.

**Fig. S3: Ub mutants of CB do not show difference in eGFP-gephyrin clustering along proximal-distal axis. (A-E’)** Morphological analysis of neurons co-transfected with eGFP-gephyin and CB1 ubiquitin mutants CB1_SH3+_ K491R/K492R or CB1_SH3+_ K491R/K492R. **(F-H)** Quantification of eGFP-gephyrin synaptic cluster density in neurons co-transfected with CB1_SH3+_ K491R/K492R or CB1_SH3-_ K431R/K432R mutants show enhanced eGFP-gephyrin cluster density (DIV 8 + 7). Scale bar 5μm. One-way ANOVA with Kruskal-Wallis non-parametric test, Dunn’s multiple comparison test p<0.0001).

**Fig. S4: eGFP-CB1_SH3-_ functions downstream of** a**2 GABA_A_Rs in newborn neurons. (A)** Morphology of adult newborn neuron expressing eGFP in WT or *Gabra2* KO background, and eGFP-CB1_SH3-_ in *Gabra2* KO background. **(B-B”)** Migration of adult newborn neurons in *Gabra2* KO background is enhanced at 14 dpi compared to WT background. WT cells catch up at 28 dpi and 42 dpi. **(C-C”)** Comparison for neuronal migration in *Gabra2* KO background show reduced migration in neurons expressing eGFP-CB1_SH3-_ in comparison to eGFP control.

